# Towards robust, unobtrusive sensing of respiration using ultra-wideband impulse radar for the care of people living with dementia

**DOI:** 10.1101/2020.12.14.422564

**Authors:** Ziwei Chen, Alan Bannon, Adrien Rapeaux, Timothy G. Constandinou

## Abstract

The unobtrusive monitoring of vital signals and behaviour can be used to gather intelligence to support the care of people living with dementia. This can provide insights into the persons wellbeing and the neurogenerative process, as well as enable them to continue to live safely at home, thereby improving their quality of life. Within this context, this study investigated the deployability of non-contact respiration rate (RR) measurement based on an Ultra-Wideband (UWB) radar System-on-Chip (SoC). An algorithm was developed to simultaneously and continuously extract the respiration signal, together with the confidence level of the respiration signal and the target position, without needing any prior calibration. The radar-measured RR results were compared to the RR results obtained from a ground truth measure based on the breathing sound, and the error rates were within 8% with a mean value of 2.4%. The target localisation results match to the radar-to-chest distances with a mean error rate of 5.4%. The tested measurement range was up to 5m. The results suggest that the algorithm could perform sufficiently well in non-contact stationary respiration rate detection.

## I. Introduction

As populations age, the demand for dementia care increases. Around 50 million people worldwide are living with dementia, with problems of memory loss, thinking and language [1], which create an urgent need to help the patients to live better and in greater safety. Respiration rate (RR) monitoring can benefit smart dementia care. Respiration rate is a key predictor of some serious clinical events, and the elderly have a higher chance of experiencing breathing difficulties due to changes in lung structure and respiratory system that come with age [2]. Specifically for dementia, there is evidence of a relationship between breathing dysrhythmias and dementia with Lewy bodies [3], and sleep-disordered breathing with Alzheimer’s Disease [4], [5]. Contactless monitoring of breathing rate could therefore help with monitoring of dementia risk factors and symptoms in the home for more effective healthcare.

Devices requiring contact with the body for measurements are less beneficial for dementia care applications. Long-term measurements are hindered by issues with patient compliance and perceived device stigmatisation [6], [7]. Dementia patients can forget to wear or charge the device, which results in higher burden compared to passive devices plugged into a wall outlet. Non-contact devices can maximise user comfort by carrying out measurements without direct interaction. Impulse Radio Ultra-Wideband (IR-UWB) radar uses GHz electromagnetic waves, and can go through obstacles including cloth and walls and reflect every object in its Field of View (FoV), making it more generally applicable. It has high resolution and robustness in the face of multi-path interference. The non-ionising waves are safe for daily use as well. Moreover, it is a platform technology that allows deriving many clinically useful measures from the same data stream, improving its scalability and affordability for home-based applications.

The literature shows that RR has been measured using UWB radar using many different algorithms. Application scenarios include in-car [8], in clinical environments [9], at home [10], [11] or portable when installed with a mobile phone [12] etc. Testing distances between object and radar are generally from 0.5 to 2 meters, except for [10], which is up to 8m. Most of the measurements require the target to be stationary, apart from [8] which takes into account the motions and hand gestures associated with driving. Systems shown in [9], [10] and [11] are relatively bulky, apart from which the system introduced in [10] is expensive. For ground truth, [9] uses manual counting, for which the accuracy depends on the observer, the presence of whom could also lead to measurement artefacts. [10] and [13] use chest sensors, giving more accurate results. The respiration signals are also used to build respiration patterns for heart rate measurement, requiring accurate measurement of the duration of each breath [13]. In addition, UWB radar can also be used for respiration detection during sleep [14], [15].

The Novelda X4M03 combines an X4 UWB radar System-on-Chip (SoC), a microcontroller and PCB antenna board. It is highly compact, has an SPI interface and is therefore highly versatile, and is low-cost [16]. The maximum radar emission is −41.3 dBm/MHz, which is within the limitations stated in 47 CFR Part 15, illustrating the amount that electronic devices can emit unintentionally [13]. It allows measurements with targets up to 9.9 metres away. These are essential features for the scalability of home-based applications. It has been shown in [16], [17] that the X4 radar SoC is capable of non-contact vital sign monitoring in humans. Our previous work demonstrated the ability to sense heart rate variability using a similar (non-contact) setup [17] and cardiovascular dynamics using a body-coupled antenna [18]. This work develops and describes a non-contact method for robustly sensing respiration rate as one component within a ‘healthy home’ platform for the care of people living with dementia. The remainder of this paper is organised as follows: Section II describes the experimental setup and data processing methods; Section III describes the experimental protocol; Section IV presents and discusses the results; and Section V draws a conclusion.

## II. Methods

### A. Experimental Setup

The experimental setup is illustrated in Fig. 1a. The radar antenna transmits UWB pulses towards the target and some pulses will be reflected by the chest wall, giving a single frame radar output as shown in Fig. 1b. Periodic chest wall movements due to respiration will then cause radar output amplitude variations. It is therefore possible to extract the RR through sampling the variations in time domain (i.e. slow-time sampling, with a configured rate of 17Hz), as displayed in Fig. 1c. Due to limited access to highly accurate respiration rate monitoring devices, a measure of ground truth respiration rate based on the breathing airflow sound is established to validate the data.

**Fig. 1.**
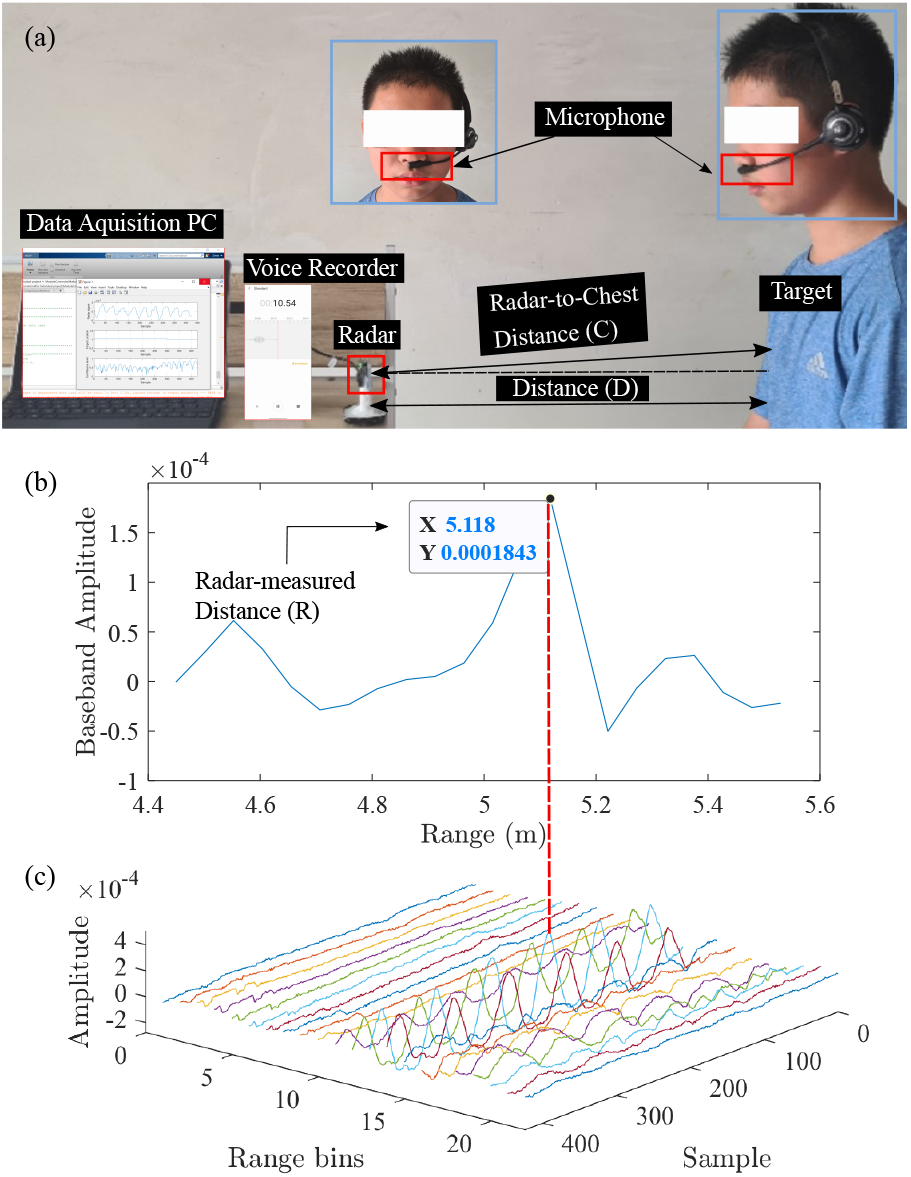
(a) Experimental setup illustration. The human target sits stationary in front of the radar with a distance (D) and a radar-chest distance (C). Results are being plotted in PC in real-time. Microphone of a Bluetooth earphone is put under the target’s nasal cavity to record the airflow sound due to breathing, which is used as a measure of ground truth. The sound are saved by the voice recorder. (b) Radar baseband output varying with range. The peak refers to the radar-measured target location. (c) Baseband amplitude variations in time. The sample axis is equivalent to a time axis. The range bin axis is equivalent to the range axis in (b), where each range bin is referred to as a fast-time (i.e. radar range sampling, with a rate of 23.328GS/s) sample point.

### B. Processing algorithm

The data processing algorithm is demonstrated in Fig. 2. A 25s’ time window is created to display the results. The range bins containing respiration signals are selected, summed and processed to give the radar signal, as displayed in Fig. 3a. The data can also be used to localise the target (Fig. 3b) and a confidence level calculation is included to check whether the radar signal is due to respiration (Fig. 3c).

**Fig. 2.**
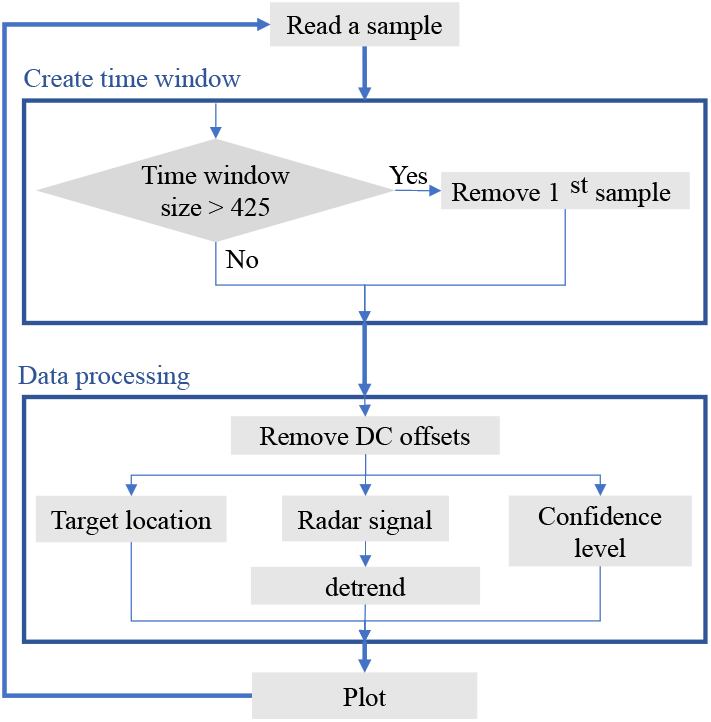
Flowchart illustrating the processing algorithm.

**Fig. 3.**
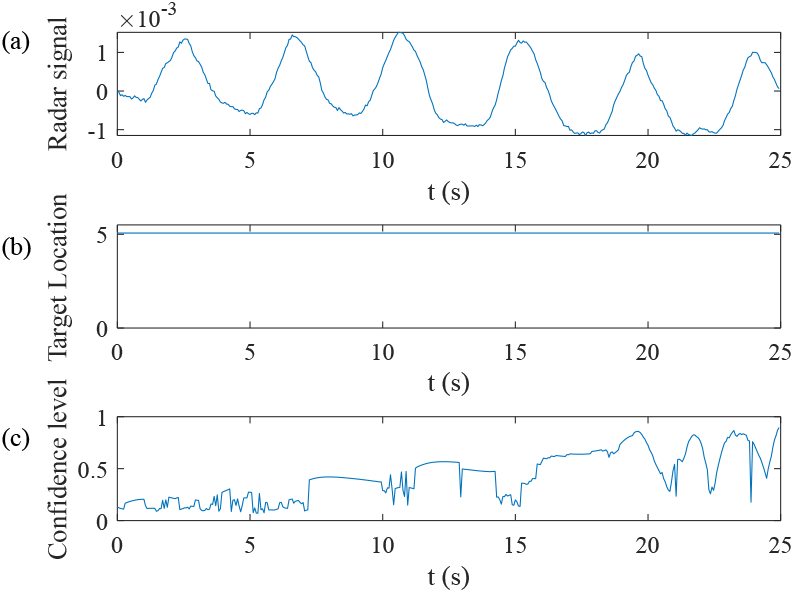
Example of output plot. (a) Radar signal. (b) Target Location. (c) Confidence level.

#### 1) Time window

In this study, a sample is received every 59ms and the window size is configured to 425 samples (i.e. 25s). The frame area is set to 1m, which concludes the human target width and ensures the real-time plot.

#### 2) Radar signal

In experimental recordings, some range bin signals are not respiration related and some may be in anti-phase to others. This algorithm takes all respiration-related range bin signals into account, thus, the first step is to select the range bin signals containing respiration information and then invert the anti-phase signals.

The method is to find out the cross-correlation between a reference range bin signal and each of the remaining range bin signals. The cross-correlation measures the similarity between two signals. If the two signals are perfectly correlated, the cross-correlation will be 1. If the cross-correlation value is higher than 0.1, the corresponding range bin signal will be selected; If the value is less than −0.1, the range bin signal will be inverted first. Finally, those signals are summed up to obtain the final radar signal. The reference range bin signal is therefore important. The range bin signal with the maximum oscillation is used, because it is the most influential signal for the final radar signal and the range bin refers to the target location. Although this will limit applications under complex circumstances, it is a reasonable choice at this stage and could be improved in the future.

The collected raw data contains strong DC offsets, which probably result from the stationary objects within the radar FoV, e.g. the stationary target and the radar hardware imperfections [13]. In order to find the range bin with maximum oscillation, the DC offsets should be removed first. Hence, the mean of each range bin signal is subtracted from it. The Matlab filtering function is not used due to its slow run time. Finally, the signal is de-trended and plotted.

#### 3) Confidence level

A measure of confidence level is established to tell if the last 25s’ data contain respiration information. Since respiration-related signals are periodic and have sinusoidal oscillations, a method is to calculate the similarity between a range bin signal and a related sample sine wave. Auto-correlation is used to build the sample sine wave. For a periodic signal, its auto-correlation sequence has the same periodic feature. Thus, it is possible to estimate the signal frequency from its auto-correlation sequence, and use this frequency to build the sample sine wave. The confidence level is then the maximum absolute cross-correlations between each range bin signal and its sample sine wave. The non-respiration signals, e.g. signals measuring the environment and body movements as shown in Fig. 4, will have high sample sine wave frequencies, and hence small cross-correlation values and low confidence levels.

**Fig. 4.**
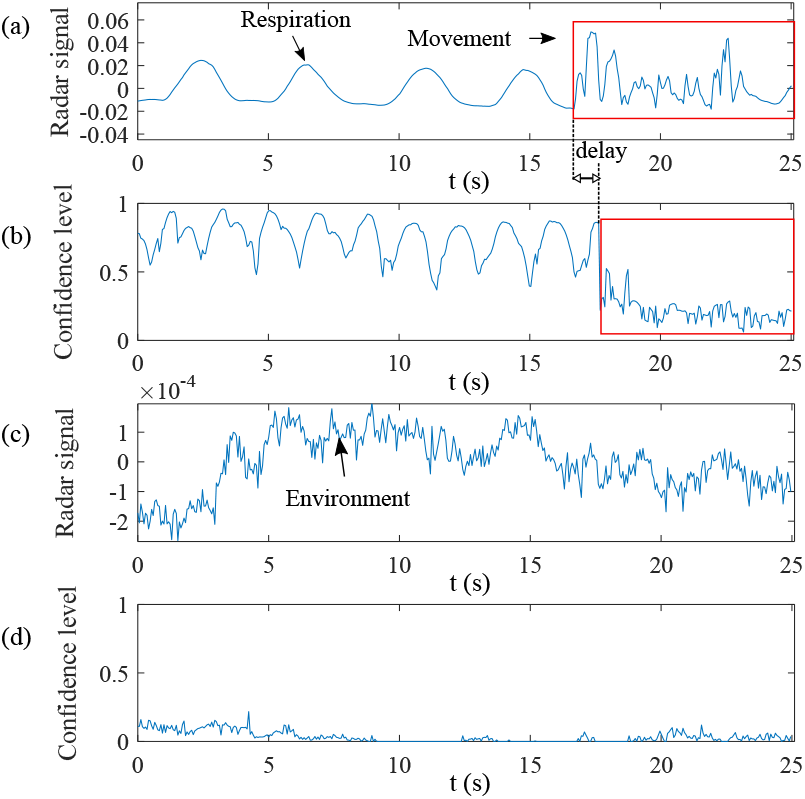
(a) Confidence level drops when target moves. (b) Low confidence level when there are no targets within the FoV.

#### 4) Target localisation

Using the respiration data instead of a single frame radar output to localise the target eliminates effects from strong stationary objects and direct path. As shown in Fig. 5, the target location obtained by a single frame radar output was altered due to the metal object, while the respiration data can reduce this effect through DC offset removal.

**Fig. 5.**
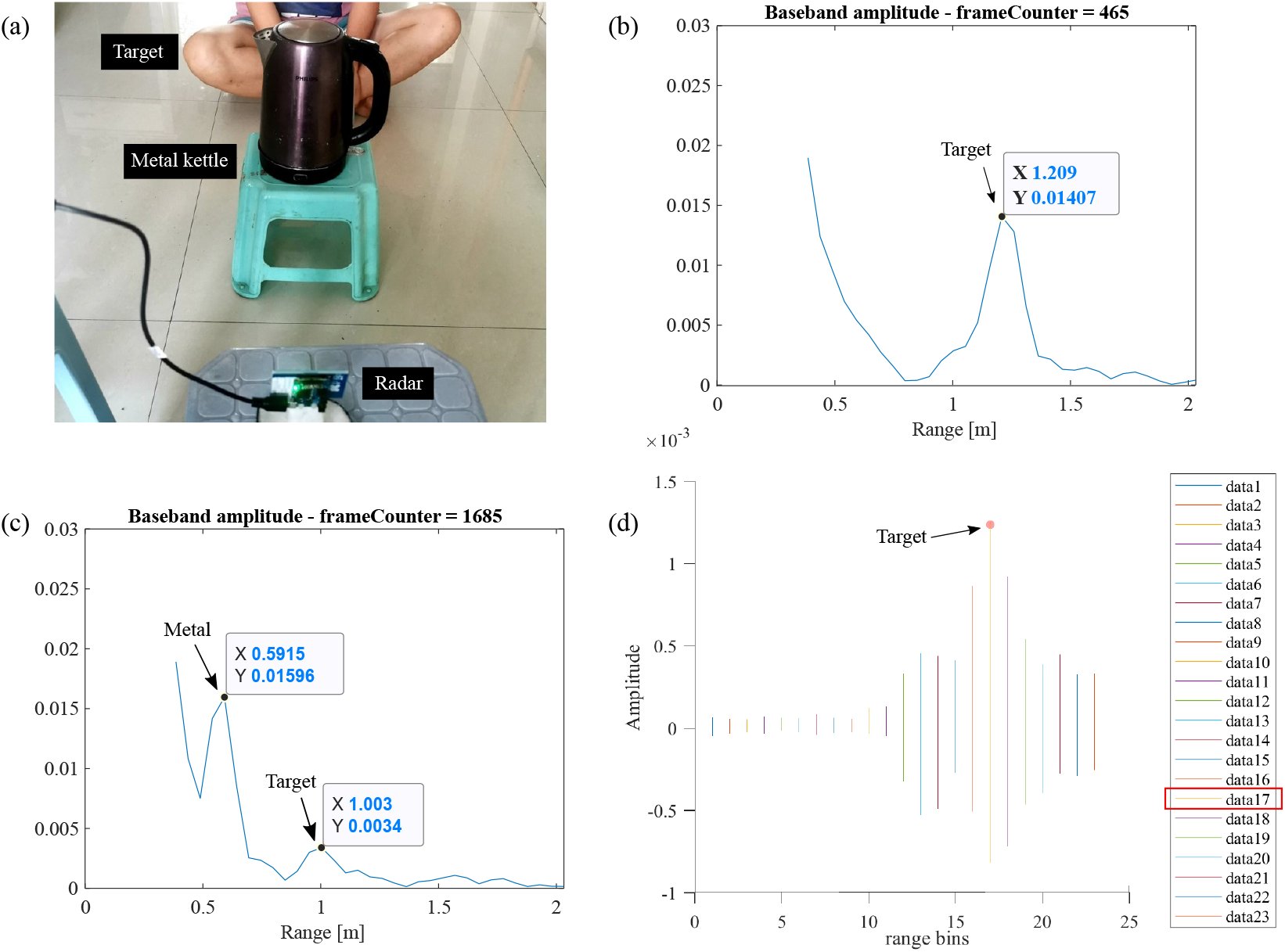
(a) Placements of radar, target, and a metal object. (b) Single-frame radar output without the kettle. The target was shown at 1.209m. (c) Single-frame radar output with the kettle. The kettle and the target were shown at 0.5915m and 1.003m respectively. (d) All bins time-domain plot. The target was shown at range bin #17, referring to 1.2088m.

## III. Experimental Protocol

In this study, we built an algorithm for UWB radar respiration rate detection. To validate the measurement, an experiment was performed on eight available targets at four different radar-target distances in a home-based environment, as demonstrated in Fig. 6. The frame range were set to 0.4-1.5m, 0.4-1.5m, 1.5-2.5m and 4.5-5.5m respectively.

**Fig. 6.**
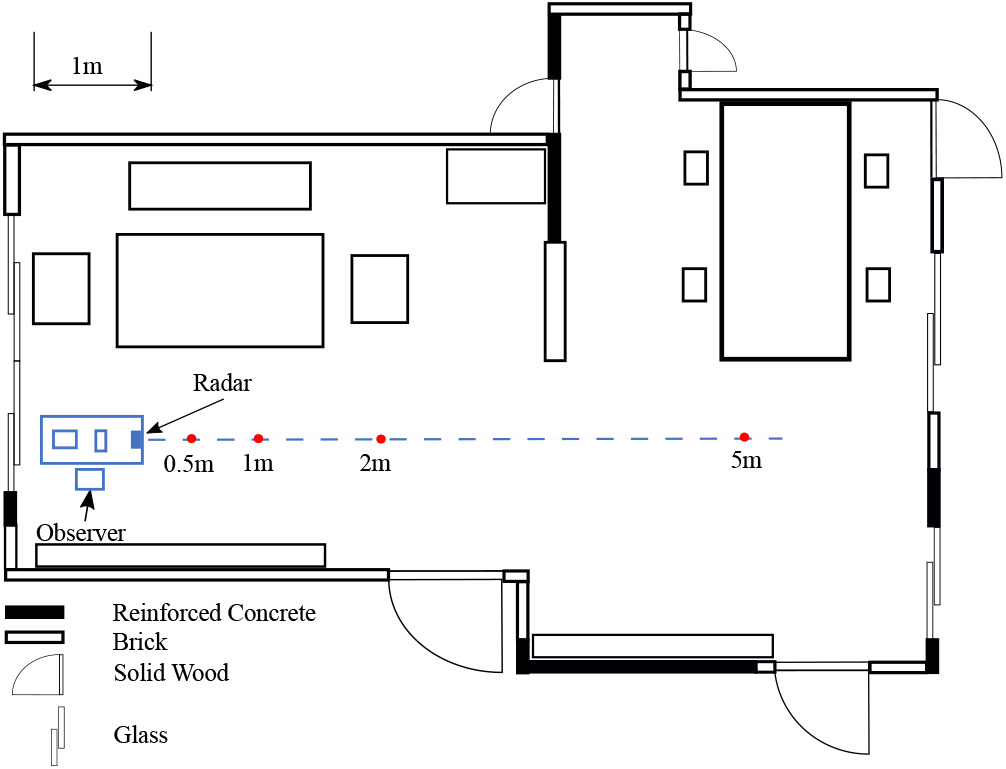
Room layout. The four possible ranges are 0.5m, 1m, 2m and 5m.

First, each target put the microphone under their nose and sat stationary in front of the radar. Then, the frame area was configured and the radar-target distance was recorded. A measurement would last for at least 25s for initialisation, and would stop when the confidence level was over at least 10%. The radar signal results were compared to the breathing sound, therefore the measurement and the voice recording were stopped at the same time. Since the plot was in real-time, the sound track end was cropped to correspond with the total sampling time.

Then, the upper envelope was estimated using spline interpolation over local maxima separated by Np samples. Default value of Np was 25,000. Because the experiment was in a home-based environment, some loud noise were however recorded. Besides, the airflow sound due to inhalation was usually much weaker compared to that of exhalation, but this is not the case when the target has a stuffy nose. Hence Np values of some tests were changed to find the best-fit envelope, as marked in Table II. An example of the comparison can be seen in Fig. 7a.

**Fig. 7.**
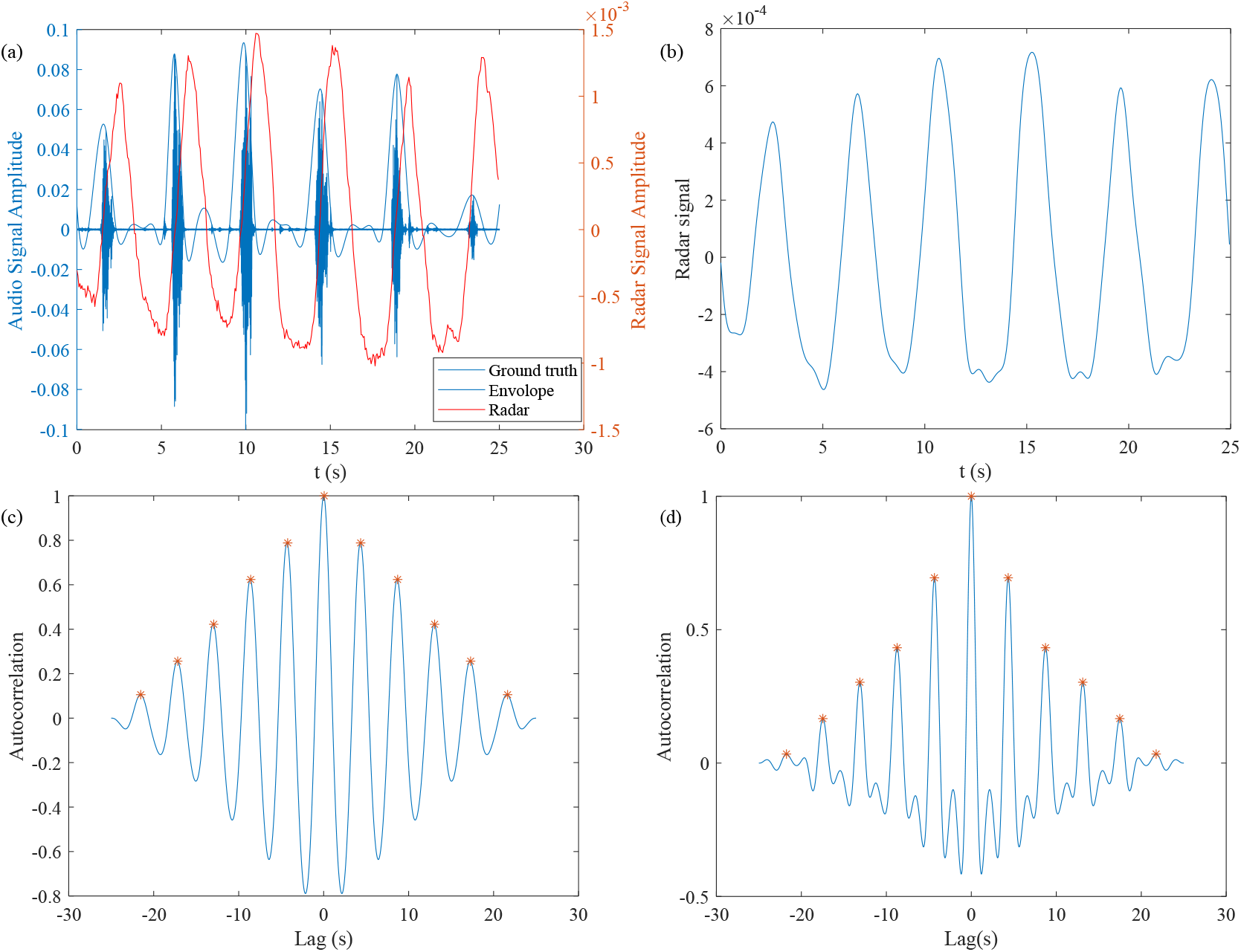
(a) Comparison of the ground truth, its upper envelope and the radar signal. (b) Out-of-band frequencies (f<0.05Hz and f>1Hz) removed. (c) Auto-correlation of (b). (d) Auto-correlation of the upper envelope.

Similar to the confidence level calculation, the RR results of two signals were calculated by the mean time differences between peaks of their auto-correlation sequences, with lags limited to ±25*s*. The out-of-band frequencies (f<0.05Hz and f>1Hz) were removed before the calculation (Fig. 7b). An example can be found in Fig. 7c and Fig. 7d.

## IV. Results & Discussion

The information of the targets and the RR results are listed in Table I and Table II, where the error rates were calculated by,

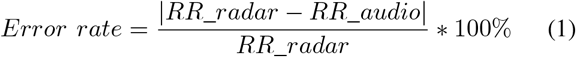

**TABLE I.** Information of targets

| Target | 1 | 2 | 3 | 4 | 5 | 6 | 7 | 8 |
| --- | --- | --- | --- | --- | --- | --- | --- | --- |
| Gender | F | M | F | F | F | M | F | M |
| Age | 47 | 49 | 39 | 23 | 12 | 25 | 11 | 12 |
| Weight(kg) | 67.8 | 70 | 50 | 65 | 52.7 | 66 | 35.5 | 57.2 |
| Height(m) | 1.65 | 1.71 | 1.63 | 1.64 | 1.62 | 1.77 | 1.45 | 1.60 |

**TABLE II.**
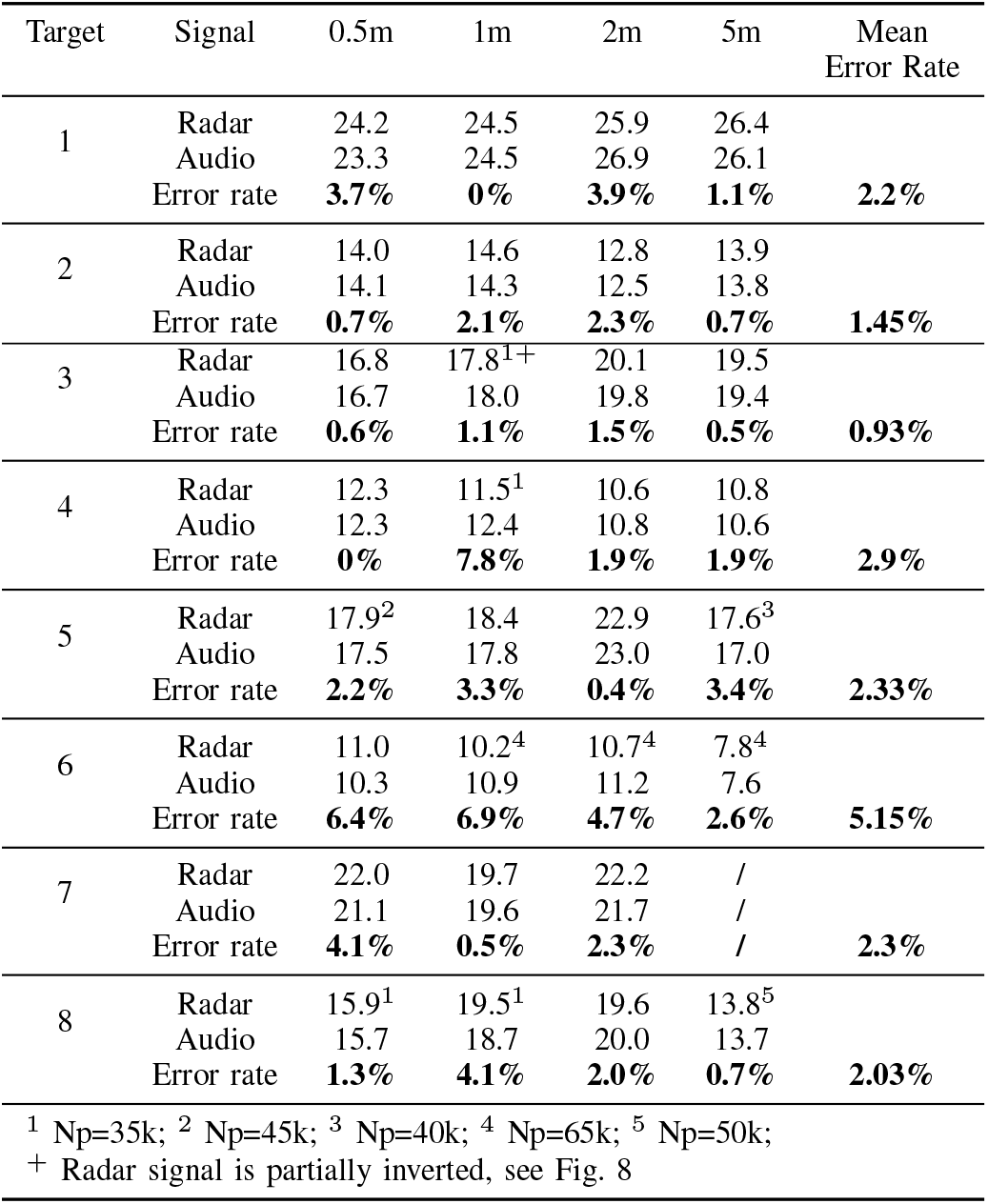
RR results (in RPM)

The error rates were within 8%, with a mean value of 2.4%, when compared to the ground truth. From this experiment, the radar signal amplitudes would decrease with distance, but the RR detection accuracy was not affected by the distance.

A limitation of the algorithm was that local maxima of the radar signal could due to either inhalation or exhalation. This was found not only from range bin to range bin, as aforementioned, but on one range bin signal, as shown in Fig. 8, where the RR result was obtained after inverting the radar signal after 6.987s. Further experiments and more strictly controlled experimental environments are required to figure out the reason. Besides, RR results of the audio signal would vary with Np. Hence, more accurate ground truth validation are recommended in future tests. In addition, home-based experimental environments introduced many reflectors that could have an impact on the measurements. It was found that the radar could also detect respiration signals when the target was sitting behind the radar. Also, there were multi-targets within the room during the experiment, e.g. the observer in Fig. 6). Those may affect the results and could also be considered and improved in the future.

**Fig. 8.**
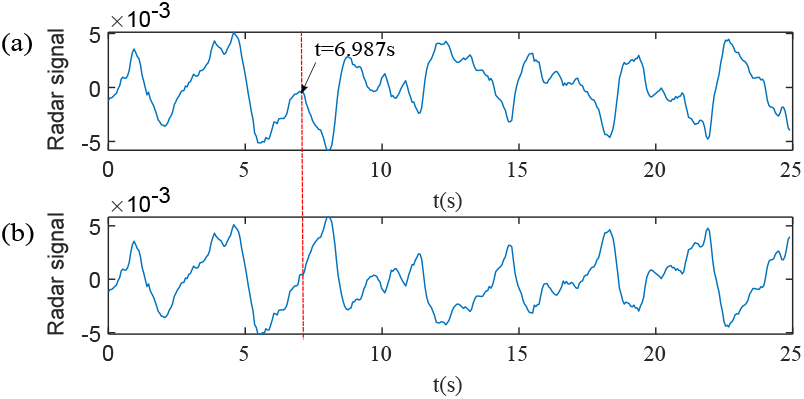
(a) Target 3’s radar signal at 1m. (b) t > 6.987s were inverted.

The confidence levels were oscillating when measuring breathing signals, such as Fig. 3c and Fig. 4b. Hence the mean and standard deviations were plotted in Fig. 9. Similar to RR results, the confidence level was not found to vary with distance. The sample sines were built with 0 phase, while the range bin signal was shifting in real-time and could have phases, hence the oscillations. A solution is to calculate the cross-correlation between each range bin and its sample sine with lags to eliminate the effect of phases. Besides, the confidence level shown in Fig. 3c was increasing, because the default values of the confidence level vector were 0 for the first 25s. In addition, delay was found as shown in Fig. 4a and Fig. 4b, which should also be improved in the future.

**Fig. 9.**
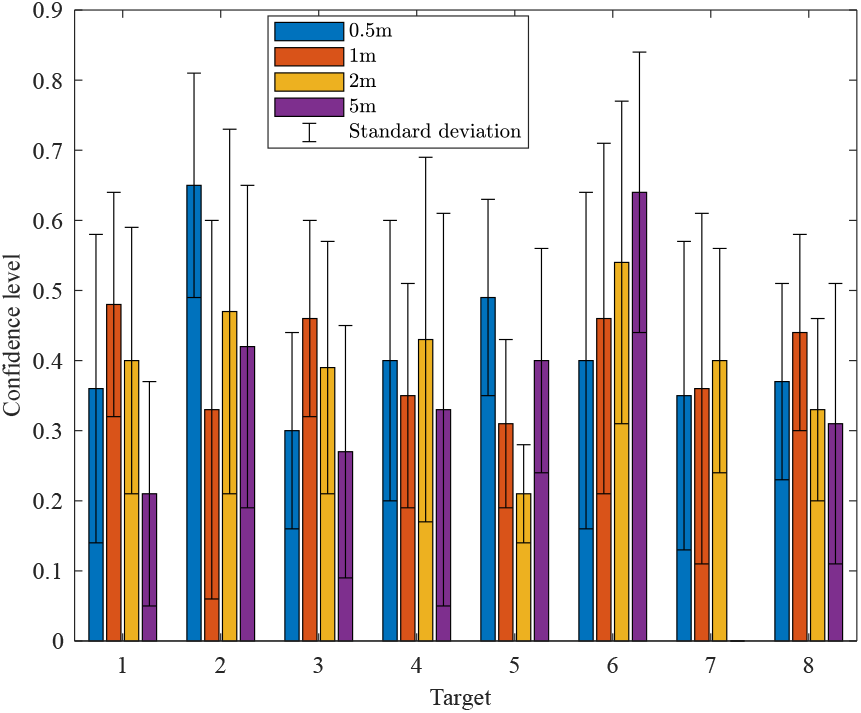
Mean and standard deviations of the last 25s’ confidence level.

The target localisation results were summarised in Fig. 10b. Each radar-measured distance value was the mean of the last 25s’ results. The error rate was calculated as,

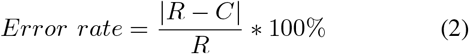

**Fig. 10.**
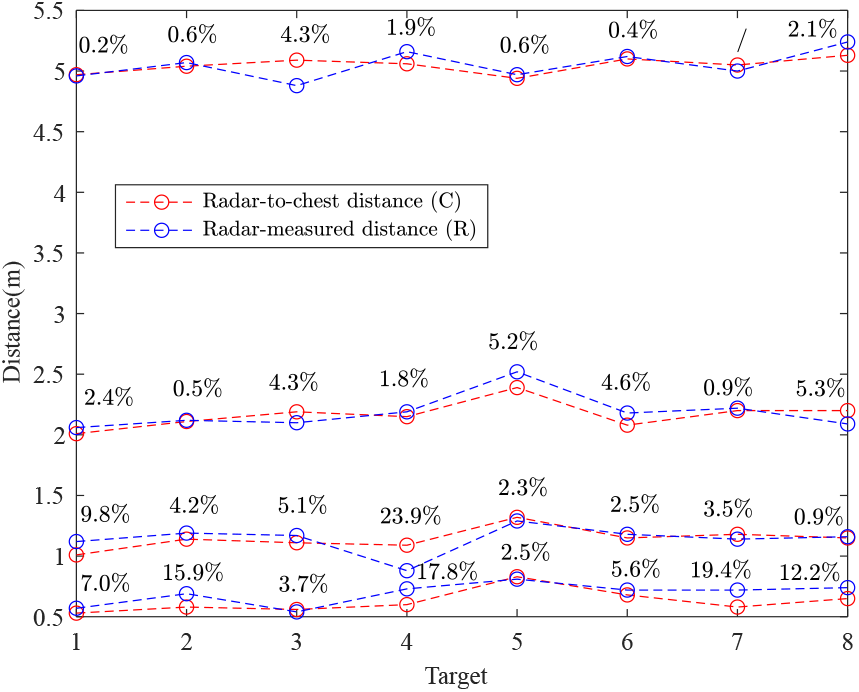
Comparison of radar-chest distance results and radar-measured distance results with error rates labelled. Mean error rates of 0.5m, 1m, 2m and 5m are 10.6%, 6.6%, 3.1% and 1.4% respectively.

The relatively high error rates at smaller distance are due to a range resolution of 0.05m. The results were compared to radar-chest distance, while the human body could overlap more than one range bin, e.g. females or heavy persons. In future validations, the radar-target distance, angles and human body width that UWB pulse can penetrate through should also be considered.

## V. Conclusion

This paper investigated the deployability of stationary RR detection using UWB radar for radar-target distance up to 5m. The radar signal, target location and confidence level of detected respiration signals were plotted in real-time. The algorithm was tested and found to perform sufficiently well for stationary RR detection. This non-contact respiration monitoring application could potentially benefit dementia care in smart home environments. It is safe for home-based environments and does not require any external sensors or devices except for the UWB radar. To build a more practical and reliable system, the movement artefacts and the inversion problems need to be investigated in future study. Besides, although the experiments were up to 5m, the radar is capable of taking measurements at distances up to 9.9 metres. This can also be tested in the future for RR monitoring.

